# Gene Therapy Rescues Cone Function in an All-cone Retina Mouse Model for Blue Cone Monochromacy with the Most Common C203R Missense Mutation

**DOI:** 10.1101/2025.09.04.674359

**Authors:** Marion E. Cahill, Kathryn Chmelik, Brooke A. Brothers, Madyson E. Ashcraft, Tongju Guan, Artjola Puja, Yinxiao Xiang, Lee M. Shaw, Jianhai Du, Wen-Tao Deng

**Affiliations:** Department of Biochemistry and Molecular Medicine, West Virginia University, Morgantown, WV 26506; Department of Ophthalmology and Visual Sciences, West Virginia University, Morgantown, WV 26506; Department of Biology, West Virginia University, Morgantown, WV 26506; Department of Microbiology, Immunology, and Cell Biology, West Virginia University, Morgantown, WV 26506

**Keywords:** blue cone monochromacy, cone opsin, cone dystrophy, AAV, gene therapy, fovea

## Abstract

Blue cone monochromacy (BCM) is an X-linked cone dystrophy characterized by loss of long- (L) and medium-wavelength (M) cone function. A common cause is the C203R missense mutation, which occurs in both *OPN1LW* and *OPN1MW*, or in hybrid *OPN1LW/OPN1MW* opsin genes. Because BCM primarily affects foveal cones, we generated *Opn1mw^C198R^/Opn1sw^-/-^/Nrl^-/-^* (*C198RAC*) mice carrying the murine equivalent of the human C203R mutation on an all-cone retinal background. C198RAC mice exhibited absent photopic ERG responses and significantly shortened cone outer segments, recapitulating foveal cone deficits in BCM. Metabolomic profiling further revealed altered retinal metabolism, including reduced cGMP and elevated oxidative stress–related metabolites.

To evaluate therapy, we delivered AAV8-Y733F expressing human L-opsin (OPN1LW) cDNA under the cone-specific PR2.1 promoter at 1 and 5 months of age. Treatment restored cone function, regenerated outer segment structures, and provided rescue for at least 5 months post-injection in both early- and late-treatment groups. These results demonstrate that densely packed cones expressing only the C198R mutant opsin remain viable targets for gene therapy.

Together, this study establishes the *C198RAC* mouse as a cone-rich model of BCM and provides compelling preclinical evidence that AAV-mediated gene augmentation can restore cone structure and function, supporting the feasibility of gene therapy for BCM.

## INTRODUCTION

Blue cone monochromacy (BCM) is an X-linked disorder characterized by complete loss or severe reduction in long-wavelength (L-) and medium-wavelength (M-) cone activity. The condition is estimated to affect approximately 1 in 100,000 individuals globally [1, 2]. From birth, individuals with BCM typically present with markedly reduced visual acuity and impaired color discrimination. Additional common features include photophobia, infantile nystagmus, and myopia [3, 4]. BCM arises from mutations within the *OPN1LW/OPN1MW* gene cluster, which encodes L- and M-opsin proteins, respectively.

These genes are arranged in a head-to-tail tandem configuration on the X chromosome, with one *OPN1LW* gene followed by one or more *OPN1MW* copies [5]. *OPN1LW* and *OPN1MW* are thought to have arisen through gene duplication, as these genes share 96% sequence homology. The sequence variation between these genes is likely to be involved in the spectral tuning of the specific opsin proteins [6]. Due to both high sequence homology and close physical proximity, these genes are prone to unequal homologous recombination and gene conversion events, resulting in the formation of hybrid opsin genes containing sequences from both *OPN1LW* and *OPN1MW* [7-9],[10-12].

One of the most prevalent causes of BCM is the Cys203Arg (C203R) missense mutation found within such hybrid opsin genes [3, 12-17]. This mutation typically arises through a two-step mechanism: initially, non-allelic homologous recombination generates a single *OPN1LW* or *OPN1LW/OPN1MW* hybrid gene; subsequently, a point mutation introduces the C203R amino acid substitution, leading to loss of L- or M-opsin function. Structural modeling based on rhodopsin suggests that C203 in cone opsins is analogous to rhodopsin C110, which forms a disulfide bond (linked to C187) essential for proper protein folding[18]. The C203R substitution is believed to disrupt this bond, resulting in misfolded opsin that fails to traffic correctly to the cone outer segment (COS). Supporting this hypothesis, *in vitro* studies show that C203R-mutant opsin is retained within the endoplasmic reticulum (ER) due to misfolding [13].

To investigate the pathogenic mechanism and therapeutic potential for BCM with the C203R mutation, we previously developed a mouse model (*Opn1mw^C198R^Opn1sw^-/^*^-^) harboring the equivalent of the human opsin mutation. These mice exhibited a phenotype consistent with human BCM patients, including absent or shortened COS and nonfunctional cones. Notably, the mutant C198R opsin was undetectable at all examined stages. Furthermore, we showed that AAV-mediated gene augmentation therapy successfully restored cone function and structure when delivered before 3 months of age. Treated cones developed well-formed outer segments and re-expressed key phototransduction proteins [19].

Despite these promising results, the retinas of *Opn1mw^C198R^Opn1sw^-/-^* mice are rod- dominant with only 3% of photoreceptor cells being cones. Unlike the human retina, where L- and M-cones are densely packed within the fovea, mouse cones are sparsely distributed across the retina. To generate a more representative model for foveal cone disorders like BCM, we crossed *Opn1mw^C198R^Opn1sw^-/-^* mice with *Nrl^-/-^* mice, which lack the neural retina leucine zipper (*Nrl*) transcription factor essential for rod differentiation. As a result, *Nrl^-/-^* mice develop all-cone retinas composed entirely of M- and S-cones [20, 21]. *Nrl^-/-^* mice have been widely adopted to study cone-specific diseases, including achromatopsia and Leber congenital amaurosis (LCA) [22-27].

Here, we demonstrated that all-cone retinas of *Opn1mw^C198R^/Opn1sw^-/-^*/*Nrl^-/-^* (*C198R-all- cone, or C198RAC*) mice lack opsin expression, exhibit abolished cone-mediated visual responses, and show a marked reduction in critical cone phototransduction proteins. These findings demonstrate that the *C198RAC* retina closely mimics the structural and functional deficits observed in the foveal cones of BCM patients with C203R mutations, offering a highly relevant model for therapeutic evaluation.

We performed gene therapy using *C198RAC* mice and showed that adeno-associated virus (AAV)-mediated gene augmentation successfully restored cone function and essential outer segment protein expression. These results support the potential of gene therapy to rescue cone function even in densely packed mutant cones expressing the C203R opsin variant, significantly advancing the feasibility of treating BCM through gene-based interventions.

## MATERIALS AND METHODS

### Animals

*C198RAC* mice were generated by crossing *Opn1mw^C198R^Opn1sw^-/-^* with *Nrl^-/-^* mice. *Opn1mw^C198R^* and *Nrl^-/-^* mice were described previously [19] [20], and the mice were genotyped by the Transnetyx outsourced genotyping service. We observed no differences in phenotype between males and females for both strains, and both sexes were used in all experiments. All animals were maintained under standard laboratory conditions (18°C-23°C, 40%-65% humidity) with food and water available ad libitum. All experimental procedures involving animals in this study were approved and conducted in strict accordance with relevant guidelines and regulations by the Institutional Animal Care and Use Committee at West Virginia University (IACUC Protocol #: 2102039943), the ARVO Statement for the Use of Animals in Ophthalmic and Vision Research, and the National Institutes of Health.

### Targeted metabolomics

Metabolites were extracted from TKO and *Nrl^-/-^* at P30 and analyzed with LC MS as described previously[28] [29]. A total of 115 metabolites were measured with optimal parameters for targeted metabolomics (Table S1). MetaboAnalyst 6.0 (https://www.metaboanalyst.ca) was used to perform multivariate analysis and generate volcano plots (P<0.05 and fold change >1.3). The ion abundance of each metabolite in TKO was divided by those from *Nrl ^-/-^* to obtain fold changes in abundance. All raw mass spectrometry data have been deposited to MassIVE (Data set identifier: MSV000097568)

### AAV Vectors

The AAV vector expressing human OPN1LW driven by PR2.1 promoter (PR2.1-hOPN1LW) was described previously [30]. This vector was packaged in AAV AAV8- Y733F and was purified according to previously published methods [31].

### Subretinal Injection

Mouse eyes were dilated using Tropi-Phen drops (Phenylephrine HCl 2.5%, Tropicamide 1%, PINE Pharmaceuticals) preceding intramuscular injection with ketamine (80 mg/kg) and xylazine (10 mg/kg) to facilitate anesthesia. Eyes to be injected were fully covered with GenTeal (0.3% Hypromellose). Under a microscope, mouse corneas were punctured with a 25-gauge needle, and 1 µL AAV particle solution (10^10^ particles/µL, 0.1% fluorescein) was injected under the retina using a blunt-end 33-gauge needle attached to a microliter syringe (Hamilton 800 Series). Following the procedure, antisedan (Orion Corporation, Espoo, Finland) was injected intraperitoneally, and neomycin/polymyxin B/dexamethasone ophthalmic ointment (Bausch & Lomb, Inc., Tampa, FL) was administered onto the cornea.

### Electroretinography

Mice were light-adapted in normal ambient white light for over 10 mintes. Eyes were dilated as described above and anesthetized using 1.5% isoflurane (2.5% oxygen) administered through a nosecone for the entire experiment. Body tempterature is maintained close to 37°C with an electronic heating pad during the procedure. Animals were placed in the center of the light dome, and ERGs were recorded from both eyes with silver wire corneal loops placed on the sclera near the limbus with 0.3% Hypromellose. The reference electrode is inserted under the scalp and a ground wire in the hind leg muscle. Animals were placed at regular room oxygen levels to awaken. The Celeris Visual Diagnostic System (Diagnosys) administered flashing light stimuli on a white light background of 30 cd/ m^2^ intensity.

### Optical Coherence Tomography

Mouse eyes were dilated before anesthetized with ketamine/xylazine as described above. GelTeal was added to each eye before imaging the retina using the Envisu Spectral Domain Optical Coherence Tomography (SD-OCT) ophthalmic imaging system (Leica Microsystems). A B-scan was acquired of size 1.96 mm^2^ centered about the optic nerve (A/B-scan = 850; B-scan = 85; frame = 1; volume = 1). Retinal layer thickness for each eye was determined by averaging measurements from 8 different points across the retinal B-scan.

### Western Blot Analysis

Retinas were extracted and flash frozen from mouse eyes promptly after CO_2_ euthanasia and homogenized in a buffer of 0.23 M sucrose, 1X protease inhibitor cocktail (Millipore Sigma), and 5mM Tris-HCl (pH 7.5) via sonication. Protein lysates were centrifuged at 13,000 rpm for 3 minutes, and 4X Laemmli Sample Buffer (BioRad) with 5% β-mercaptoethanol was added to a 1X concentration to the samples. 100 µg protein per sample was run in a 10% Mini-PROTEAN TGX gel (BioRad) by electrophoresis. The blot was transferred to a Low Fluorescence PVDF membrane (BioRad), blocked with Intercept Blocking Buffer (LI-COR), and probed with TUB4A (MilliporeSigma cat. T5168; 1:4,000 dilution) and OPN1LW/MW (MilliporeSigma cat. AB5405; 1:1,000 dilution) primary, followed by anti-rabbit-680 (LI-COR cat. 68023; 1:20,000 dilution) and anti-mouse (LI-COR cat. 32212; 1:20,000 dilution) secondary antibodies. The blot was imaged using the Odyssey Infrared Imager (LI-COR).

### Preparation of Retinal Cross Sections and Immunohistochemistry

Mouse eyes were enucleated promptly after euthanasia, and a large hole was created along the corneal ridge using a 20-gauge needle. The eyes were incubated at room temperature in 4% paraformaldehyde (1X PBS) prior to and following the surgical removal of the cornea, for a total of 2 hours. Eyes were transferred to 20% sucrose (1X PBS) overnight at 4°C, and then moved to 10% sucrose, 50% Tissue-Tek O.C.T. compound (Sakura Finetek USA) (0.5X PBS) for 1 hour at 4°C after removing the lens. Eyes were flash frozen in O.C.T. blocks, which were then sectioned (16-µm slices) using the MES1000+ Cryostat and placed on Superfrost Plus microscope slides (Thermo Fisher) for staining. Slides were rinsed briefly in 1X PBS, blocked with 1% BSA (1X PBS), and stained with biotinylated PNA (Vector Laboratories cat. B1075; 1:5,000 dilution) and OPN1LW/MW (Kerafast cat. EJH006; 1:1,000 dilution) primary, followed by fluorescein-avidin-D-488 (Vector Laboratories cat. A2001; 1:500 dilution), anti-chicken-594 (Thermo Fisher cat. A32759; 1:500 dilution), and DAPI (Thermo Fisher cat. D1306; 1:1,000 dilution) secondary staining. Glass coverslips were mounted with Prolong Gold Antifade mountant (Thermo Fisher) and cured overnight. Images were captured on the Nikon C2 confocal microscope and analyzed in ImageJ and Photoshop software.

### Transmission Electron Microscopy of Cones

Methodology for the collection and preparation of mouse eyes for TEM imaging are previously described (PMID: 34929159, currently ref 38). Briefly, eyes were enucleated promptly after euthanasia, and a large slit was cut across the cornea. Eyes were fixed in 2% PFA and 2.5% glutaraldehyde (100 mM cacodylate, 1X PBS) before removal of the cornea and lens. Eyes remained in fixative for at least 2 days before staining the tissue with 2% osmium tetroxide (0.1 M cacodylate buffer, 1% uranyl acetate), dehydration, and embedding in Polybed 812 resin (Polysciences Inc.). Mounted tissue was stained with 3% Reynold’s lead citrate, and the JEOL JEM-1010 transmission electron microscope was used to collect images. Images were further processed in Photoshop.

### Statistical Analysis

GraphPad Prism was used to perform unpaired student’s *t-*test on data of 2 groups, 1-way ANOVA with Tukey’s post-hoc test for 3 or more groups, and 2-way ANOVA with Tukey’s post-hoc test for multiple comparisons as appropriate. Figure legends specify replicate size. All error bars represent the standard deviation from the mean. Significance is indicated as \**p* ≤ 0.05, \*\**p* < 0.002, or *\*\*\*p* < 0.001, *\*\*\*\*p* < 0.0001.

## Results

### Characterization of retinal of *Opn1mw^C198R^/Opn1sw^-/-^/Nrl^-/-^* (*C198RAC*) mice

Retinal degeneration in *Nrl^-/-^* mice begins within the first four months and slows over time, stabilizing by around 10 months of age. Their retinas show disorganization marked by rosettes and ring-shaped ONL deformities, which are most prominent in young mice and diminish with age [20, 32]. We characterized the retinal morphology of *C198RAC* mice by comparing retinal thickness with *Nrl^-/-^* mice. Optical coherence tomography (OCT) measurements of the outer nuclear layer (ONL) thickness at P30, P120, and P300 confirmed previous studies that *Nrl^-/-^* retinas degenerated initially, and then became stabilized. Compared to *Nrl^-/-^* mice, the retinas of *C198RAC* mice are slightly thicker, although the difference is not statistically significant. *C198RAC* retinas also experience degeneration initially, but at a lesser extent than *Nrl^-/-^* mice (Fig. 1).

**Figure 1.**
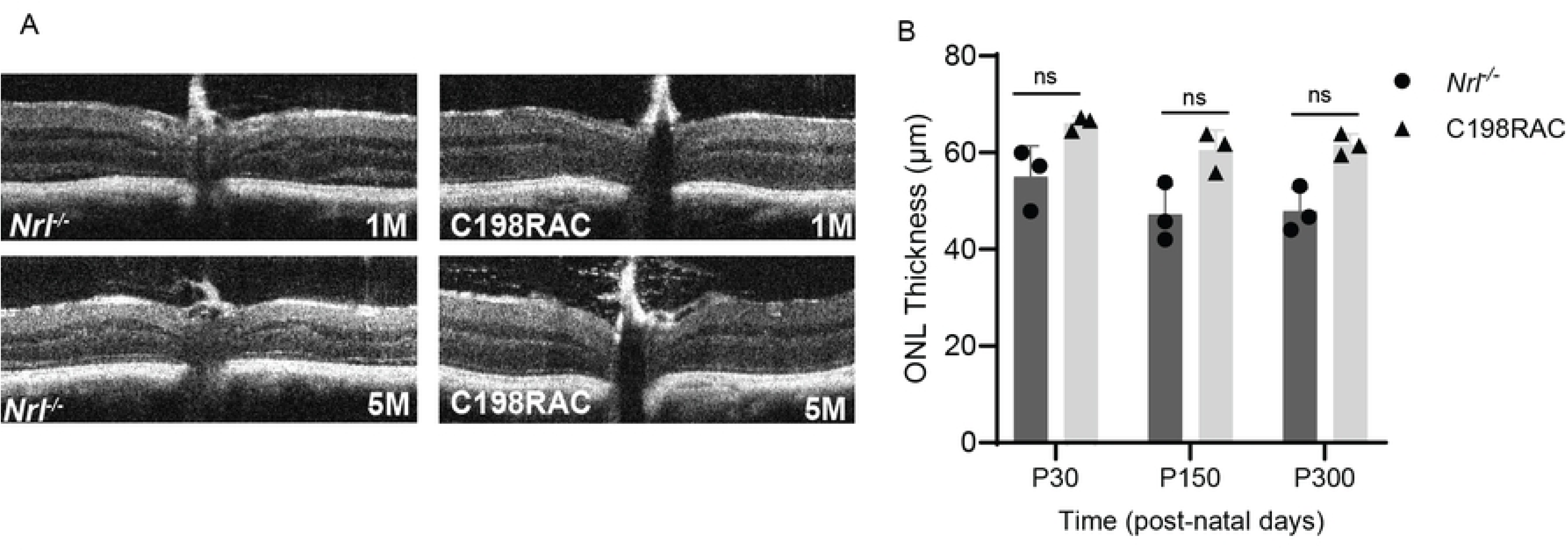
Characterization of the retinal structure of *C198RAC* mice. (A) Representative Optical Coherence Tomography (OCT) images of 1- and 5-month-old *C198RAC* and *Nrl^-/-^* mice. (B) OCT measurements of outer nuclear layer (ONL) thickness from *C198RAC* and *Nrl^-/-^* retinas at ages of postal natal day P30, P150, and P300 (n=3 eyes from different mice/group). Ns = not significant.

### C198R mutation alters retinal metabolism

To assess how the C198R mutation in *Nrl^-/-^* background affects retinal metabolism in all-cone mice, we performed targeted metabolomics using liquid chromatography mass spectrometry (LC MS) on retinas from C198RAC and *Nrl^-/-^* mice at P30. PLS-DA analysis revealed distinct retinal metabolomic profiles between C198RAC and *Nrl^-/-^* mice, indicating that the opsin mutation has a pronounced impact on retinal metabolism (Fig. 2A). Among the 115 metabolites measured, five were significantly altered in C198RAC retinas. Short-chain acylcarnitines (C4:0, isoC4:0), pantothenic acid, and ophthalmic acid were elevated, whereas cGMP—a key photoreceptor outer segment messenger for phototransduction—was markedly reduced (Fig. 2B).

**Figure 2.**
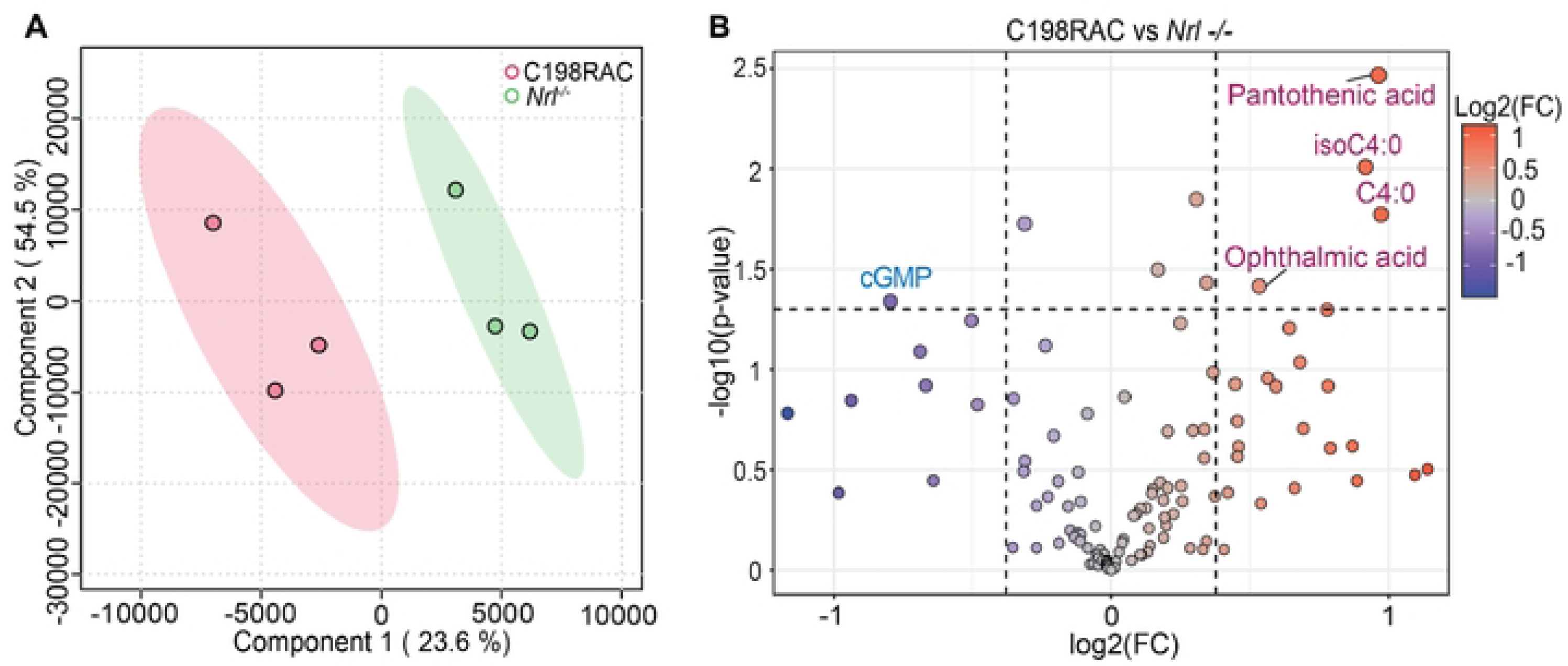
Impact of C198R mutation on retinal metabolism in *Nrl -/-* background mice. (A) Partial least squares discriminant analysis (PLS-DA) of retinal metabolomics done at one months of age (P30) distinguishes C198RAC mice from *Nrl -/-*. (B) Volcano plot showing differentially expressed metabolites between C198RAC and *Nrl-/-* mice. Significantly upregulated and downregulated metabolites are shown in red and blue, respectively (*p < 0.05, Fold Change > 1.3, N=3). Metabolites are abbreviated as follows: isobutyryl-L-carnitine (isoC4:0); butyryl-L-carnitine (C4:0); cyclic guanosine monophosphate (cGMP).

### Gene augmentation therapy showed long-term functional rescue in *C198RAC* mice treated at 1 month of age

ERG responses in *Nrl^-/-^* mice decline with age, becoming significantly reduced by 5 months and reaching about one-third of the 4-week-old amplitude by 7 months. Beyond this point, ERG function stabilizes and remains unchanged up to 12 months [33]. *C198RAC* mice showed no photopic ERG responses at one month of age (Fig. S1). We conducted gene therapy in *C198RAC* mice at one month of age (1M) using subretinal injection of AAV8-Y733F carrying human L-opsin (*OPN1LW*) cDNA under the cone-specific PR2.1 promoter. Cone function was analyzed at 1 month (1M+1M) and 5 months (1M+5M) post-injection.

**Figure S1.** A representative ERG waveforms of 1 month old C198RAC mouse and a age-matched *Nrl^-/-^* control.

At 1M+1M, treated *C198RAC* eyes showed a mean photopic b-wave amplitude of 199 ± 54 µV (n=5), significantly higher than uninjected contralateral eyes, which showed no measurable response (P < 0.0001). The rescue is ∼49% of age-matched *Nrl^-/-^* controls (408 ± 40 µV, n=5/group, P < 0.0001) at light intensity of 120 cd·s/m² (Fig. 3A & 3B). By five months post-injection (1M+5M), b-wave amplitudes in treated eyes declined to 134 ± 11 µV, which is ∼67% of 1M+1M treated eyes. Rescue at both 1M+1M and 1M+5M timepoints are still significantly lower than age-matched 2M and 6M *Nrl^-/-^* controls (408 ± 40 µV and 193 ± 26 µV, respectively; n=5/group, P < 0.0001), with 6M *Nrl^-/-^* mice exhibiting ∼50% of 2M *Nrl^-/-^* ERG levels (Fig. 3C & 3D). These results demonstrated that while treated *C198RAC* eyes show some functional decline over time, it is less pronounced than the natural age-related decline seen in *Nrl^-/-^* mice.

**Figure 3.**
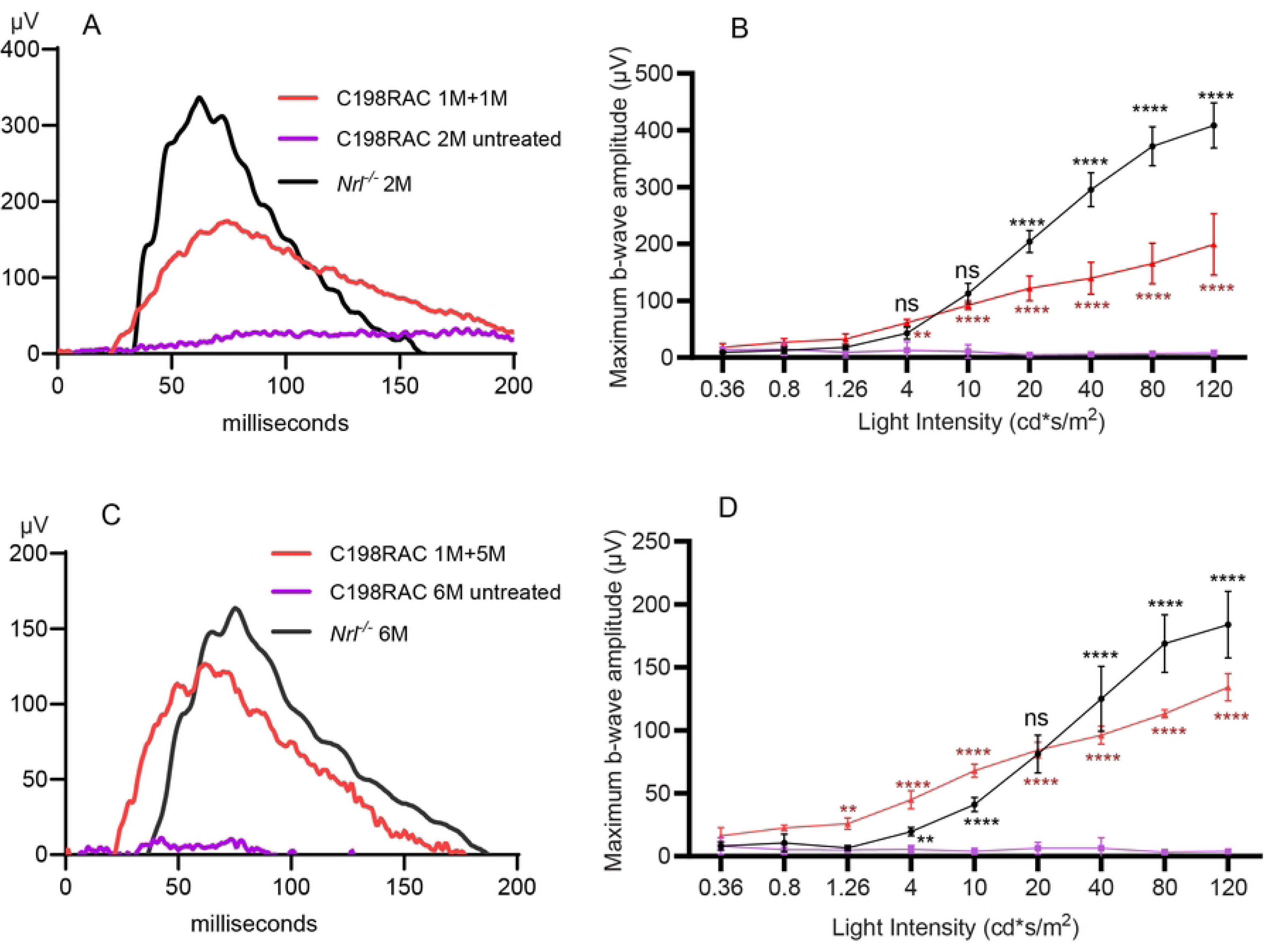
Gene therapy rescued cone function in *C198RAC* mice treated at one month of age. (A, C) Representative ERG waveforms from treated 1M+1M (A) and 1M+5M (C) *C198RAC* eyes (red line), with contralateral untreated eyes (magenta line) and age-matched *Nrl^-/-^* controls (black line), at a white light intensity of 120 cd*s/m^2^. (B, D) Photopic b-wave responses with increasing light intensities from light-adapted *C198RAC* mice treated at 1 month of age and analyzed at 1 month (B) and 5 months (D) post-injection. Untreated contralateral eyes were used as negative controls, and age-matched *Nrl^-/-^* mice were used as positive controls. Each data point represents the mean ± SD of b-wave amplitudes recorded for each group at the indicated flash intensity (n=5 for each group). Red asterisks (*) indicate statistical analysis between treated vs. untreated eyes, and black asterisks indicate statistical analysis between treated vs. *Nrl^-/-^* controls. ns = not significant, **P < 0.01, ****P < 0.0001.

### Long-term functional rescue was achieved in *C198RAC* mice following treatment at 5 months

To assess whether gene therapy remains effective at later stages, *C198RAC* mice were treated at 5 months of age using the same AAV vector, and efficacy was evaluated at 1 (5M +1M) and 5 months (5M+5M) post-injection. Treatment at this age still restored cone function, with rescue lasting at least 5 months. In 5M+1M *C198RAC* mice, the average photopic b-wave amplitude was 107 ± 46 µV at 120 cd·s/m², not significantly lower than age-matched 6-month-old *Nrl^-/-^* controls (n=6/group, P > 0.05) (Fig. 4A, 4B). At 5M+5M, amplitudes of treated mice reduced to 66 ± 6 µV, significantly lower than age-matched *Nrl^-/-^* controls (n=5/group, P < 0.0001) (Fig. 4C, 4D).

**Figure 4.**
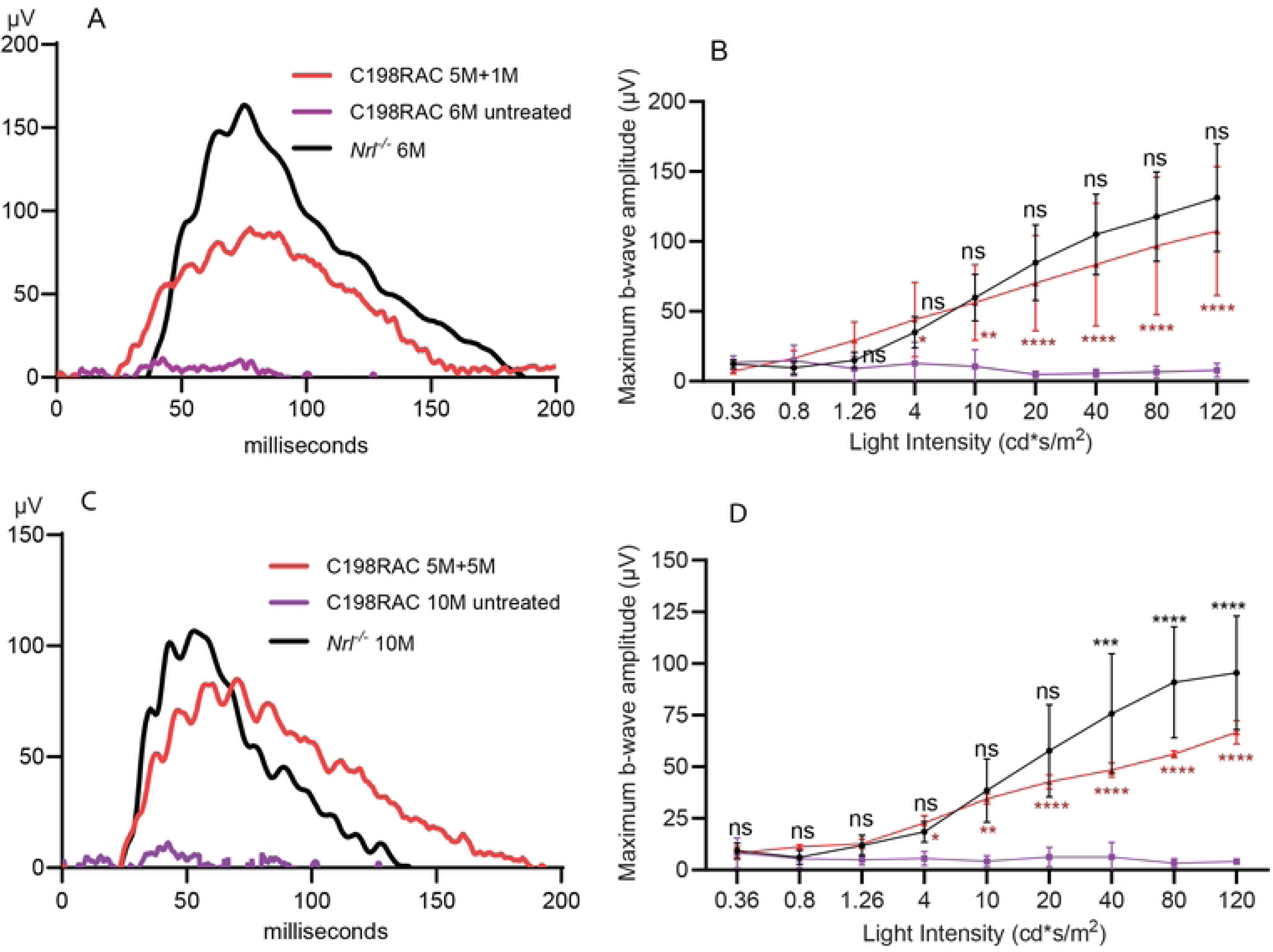
Gene therapy rescued cone function in *C198RAC* mice treated at five months of age. (A, C) Representative ERG waveforms from treated 5M+1M (A) and 5M+5M (C) *C198RAC* eyes (red line), with contralateral untreated eyes (magenta line) and age-matched *Nrl^-/-^* controls (black line), at a light intensity of 120 cd*s/m^2^. (B, D) Photopic b-wave responses with increasing light intensities from light-adapted *C198RAC* mice treated at 5 months of age and analyzed at 1 month (B) and 5 months (D) post-injection. Untreated contralateral eyes were used as negative controls, and age-matched *Nrl^-/-^* mice as positive controls. Each data point represents the mean ± SD of b-wave amplitudes recorded for each group at the indicated flash intensity (n=6 for 5M+1M group, and n=5 for 5M+5M group). Red asterisks (*) indicate statistical analysis between treated vs. untreated eyes, and black asterisks indicate statistical analysis between treated vs. *Nrl^-/-^* controls. ns = not significant, *P < 0.05, **P < 0.01, ****P < 0.0001.

### Gene therapy restored opsin expression in COS

We analyzed AAV-mediated L-opsin expression by IHC in 1M+5M and 5M+5M treated *C198RAC* eyes and demonstrated that AAV-delivered L-opsin colocalized with cone marker peanut agglutinin (PNA) (Fig. 5). In contrast, untreated contralateral eyes showed strong PNA staining, indicating viable cones, but no M-opsin staining, consistent with our previous observation that C198R mutant opsin is efficiently degraded [19]. Western blot analysis of 1M+5M and 5M+5M *C198RAC* retinas confirmed robust L-opsin expression in treated retinas, while untreated eyes lacked detectable M-opsin (Fig. 6). The L-opsin levels in treated *C198RAC* eyes were drastically higher than endogenous M-opsin levels in *Nrl^-/-^* retinas. This difference is because *Nrl^-/-^* retinas contain a higher proportion of S-cones than M-cones. Previous studies showed that the PR2.1 promoter drives transgene expression in mouse S-cones [34, 35]. Therefore, the higher L-opsin levels in treated eyes is resulted from AAV-mediated L-opsin transducing both M- and S-cones in the *C198RAC* retinas.

**Figure 5.**
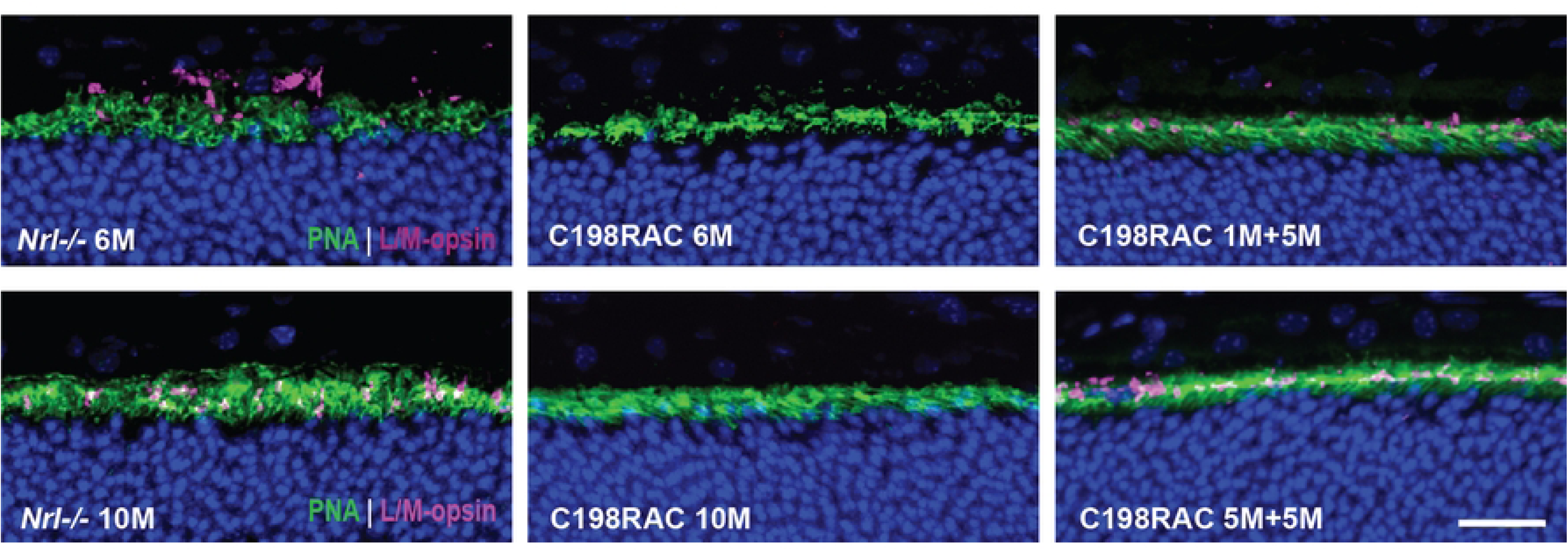
Gene therapy restored opsin expression in COS of treated *C198RAC* mice. (Top panels) Immunohistochemistry of representative retinal cross sections from a *C198RAC* eye treated at 1 month of age and collected at 5 months post-injection (1M+5M; right), with an age-matched untreated *C198RAC* (middle) and *Nrl^-/-^* control (left). (Bottom panel) Representative retinal cross sections from a *C198RAC* eye treated at 5 months of age and collected at 5 months post-injection (5M+5M; right), with an age-matched untreated *C198RAC* (middle) and *Nrl^-/-^* control (left). Staining was performed with antibody against L/M-opsin (magenta) together with PNA (green). Scale bar = 20 µm.

**Figure 6.**
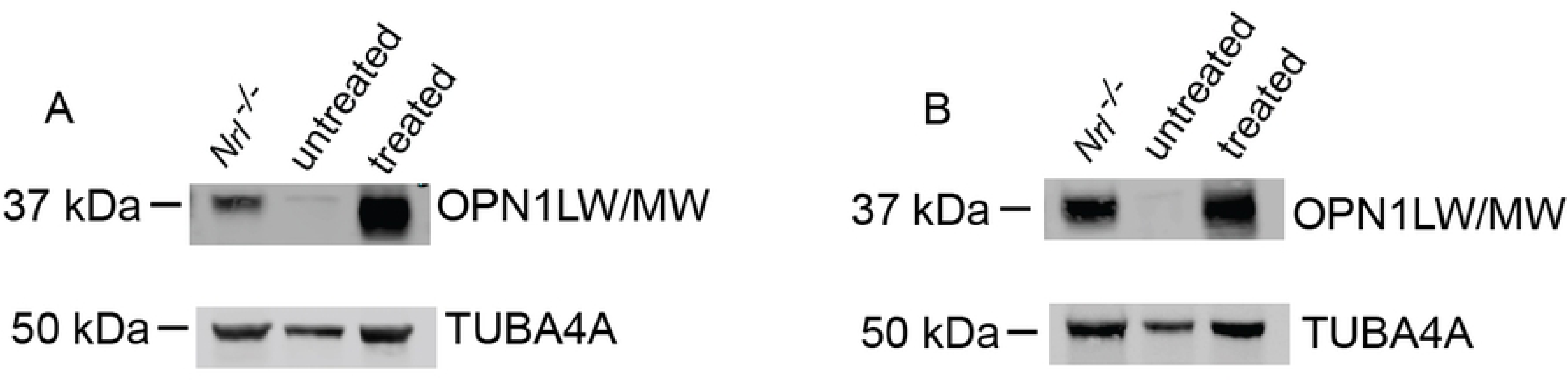
Western blot analysis of total retinal lysate from (A) 1M+5M treated *C198RAC*, age-matched untreated *C198RAC* control, and *Nrl^-/-^* eyes and (B) 5M+5M treated *C198RAC*, age-matched untreated *C198RAC* control, and *Nrl^-/-^* eyes, probed for L/M- opsin and TUBA4A as a loading control.

### Gene therapy regenerated cone outer segments in treated *C198RAC* mice

We previously showed that *Opn1mw^C198R^/Opn1sw^-/-^* mice have severely shortened or absent cone outer segments (COS), and that gene augmentation regenerated COS structure [19]. To assess whether functional rescue in treated *C198RAC* eyes also led to COS regeneration, we used transmission electron microscopy (TEM) to assess COS structure in treated *C198RAC* eyes. Untreated eyes showed only residual membrane structures adjacent to the RPE, lacking the organized, stacked morphology typical of COS (Fig. 7A, 7D). In contrast, we observed examples of well-organized COS in treated eyes (Fig. 7B, 7E), though they were less abundant than in *Nrl^-/-^* controls (Fig. 7C, 7F).

**Figure 7.**
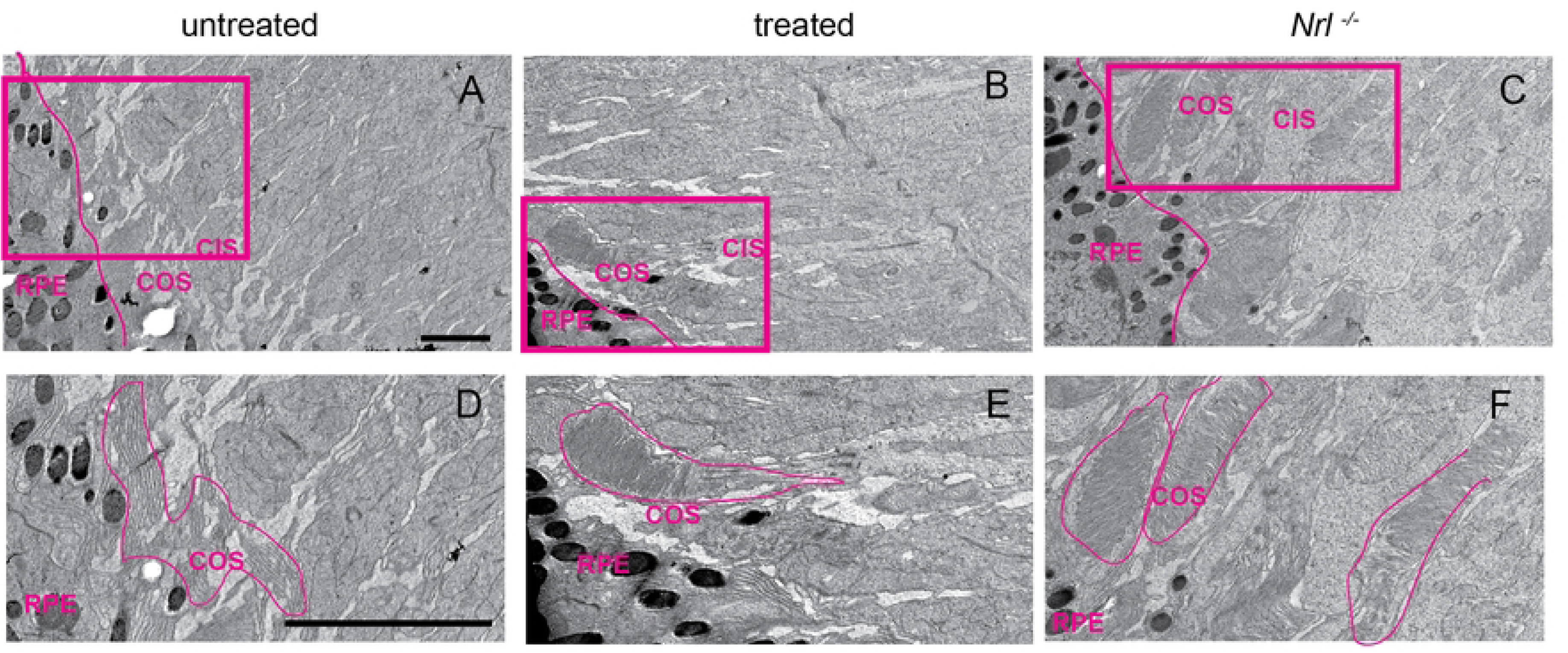
Gene therapy rescued cone outer segment structure in *C198RAC* retinas. (A, D) Representative TEM images of a 1-month-old untreated *C198RAC* retina showing residual fragmented COS membrane structure. (B, E) Representative 1M+1M treated *C198RAC* retina, showing examples of well-organized COS structures. (C, F) Representative 1-month-old *Nrl^-/-^* retina showing normal COS structure. Panels D, E, and F are zoomed-in areas of the boxed regions from A, B, and C, respectively. Magenta lines were manually drawn between RPE and COS in A, B, and C, and around COS structures in D, E, and F. Both scale bars = 2 μm.

## Discussion

In this study, we demonstrate that AAV-mediated gene augmentation therapy effectively restores cone function and structure in a cone-rich mouse model of blue cone monochromacy (BCM) carrying the C198R missense mutation, homologous to the human C203R mutation commonly associated with BCM. By leveraging the *Nrl^-/-^* background to generate all-cone retinas, the *C198RAC* model allowed us to closely mimic the high-density cone environment of the human fovea, offering a valuable platform for evaluating therapeutic strategies.

The metabolomic analysis of one-month-old *C198RAC* retinas revealed several significant alterations, providing insight into the pathophysiology of this BCM model. Most notably, a substantial reduction in cGMP levels was observed. cGMP acts as a crucial second messenger in the phototransduction cascade, binding to cyclic nucleotide-gated channels (CNGC) to sustain the photoreceptor dark current [36, 37]. This reduction in cGMP likely disrupts cone phototransduction signaling and directly contributes to the absence of photopic ERG responses observed in C198RAC mice, establishing a mechanistic link between metabolic dysfunction and the functional phenotype. Additionally, key metabolites linked to oxidative stress and metabolic dysregulation – both hallmark features of retinal degeneration – were elevated, including ophthalmic acid, short-chain acylcarnitines, and pantothenic acid. These changes suggest that the C198R mutation triggers broader metabolic disruption beyond the phototransduction defect. Ophthalmic acid, a tripeptide analog of glutathione, is known to increase in response to oxidative stress [38-40], indicating that *C198RAC* retinas experience elevated oxidative burden despite their young age. The accumulation of acylcarnitines, including butyryl-L-carnitine (C4:0) and isobutyryl-L-carnitine (isoC4:0) suggests impaired fatty acid and amino acid metabolism in *C198RAC* retinas. The elevation of pantothenic acid (vitamin B5), a precursor of coenzyme A (CoA), that is critical for the synthesis of acyl CoAs, further supports the impaired lipid metabolism [41], and bioenergetics in cones [42]. Taken together, these findings demonstrate that the C198R mutation triggers a cascade of metabolic dysfunction that may contribute to the long-term viability of cone photoreceptors in this model.

Our findings show that subretinal delivery of AAV8-Y733F encoding human L-opsin under the cone-specific PR2.1 promoter leads to robust and sustained rescue of cone-mediated visual function when administered at both one month and five months of age. This was supported by restored photopic ERG responses and regeneration of COS confirmed by TEM. Notably, these effects persisted for at least five months post-treatment, indicating long-term therapeutic benefit.

To assess the relationship between genotype and disease severity in BCM, cone photoreceptor structure was previously compared in patients with the two most common mutations: (A) large deletions and (B) the C203R missense mutation in the *OPN1LW/OPN1MW* gene cluster. Notably, patients with the C203R mutation were reported to retain foveal cone outer nuclear layer (ONL) thickness for decades and exhibited slower degeneration of inner and outer segments compared to those with deletion mutations [43]. Because the C203R mutant protein is misfolded, it was initially suspected to be toxic to cones, similar to certain rhodopsin missense mutations that cause dominant retinitis pigmentosa. Thus, the relatively mild phenotype in C203R patients was unexpected. Consistent with these clinical findings, our data show that cones in the *C198R* and *C198RAC* mouse models do not degenerate more rapidly than those in the deletion model [19], reinforcing the relevance of this model for therapeutic testing.

Importantly, we extended our findings to show that gene therapy remains effective when administered at a later stage. Treatment at five months of age also restored cone function and COS protein expression, with functional rescue maintained over five additional months. Although the magnitude of ERG responses was modestly reduced compared to earlier treatment, the rescue effect was sustained, indicating that a therapeutic window exists beyond early postnatal stages.

Our study offers compelling evidence that BCM-associated structural and functional cone deficits caused by C203R mutations can be reversed through gene supplementation, even in a densely packed, cone-dominant retina. This contrasts with previous assumptions that misfolded opsins necessarily lead to rapid cone degeneration. Instead, our data suggest that cones expressing C198R opsin remain viable for extended periods despite a lack of function and atrophied COS.

Nevertheless, some limitations remain. While *Nrl^-/-^* mice provide a valuable all-cone model, they lack the foveal architecture of the human retina. Further validation in large animal models with developed foveae, such as non-human primates, will be helpful to confirm the translational potential of AAV gene therapy for cone dystrophies. Additionally, the long-term durability of rescue beyond five months and the functional integration of restored cones into complex visual circuits require further investigation.

In conclusion, our results establish that gene augmentation therapy targeting cone opsin deficiency can effectively restore cone structure and function in a clinically relevant BCM model. The success of both early and delayed treatment provides critical support for developing gene-based interventions for BCM patients, including those beyond infancy, and highlights the therapeutic potential of targeting misfolded opsin mutations with gene supplementation strategies.

## ACKNOWLEDGMENTS

We thank Robert J. Barbera and Emily R. Sechrest for technical support.

## Data Availability

All relevant data are within the manuscript and its Supporting Information files.

